# Natural genetic variation in *GLK1*-mediated photosynthetic acclimation in response to light

**DOI:** 10.1101/2023.10.28.564491

**Authors:** Jose M Muino, Christopher Großmann, Tatjana Kleine, Kerstin Kaufmann

## Abstract

GOLDEN-like (GLK) transcription factors are central regulators of chloroplast biogenesis in Arabidopsis and other species. Findings from Arabidopsis show that these factors also contribute to photosynthetic acclimation, e.g. to variation in light intensity, and are controlled by retrograde signals emanating from the chloroplast. However, the natural variation of GLK1-centered gene-regulatory networks is largely unexplored. By evaluating the activities of GLK1 target genes and GLK1 itself in vegetative leaves of natural Arabidopsis accessions grown under standard conditions, we uncovered a remarkable variation in the activity of GLK1 centered regulatory networks. This is linked with ecogeographic origin of the accessions, and can be associated with a complex genetic variation across loci acting in different functional pathways, including photosynthesis, ROS and brassinosteroid pathways. Our results identify candidate upstream regulators that contribute to GLK1 activity in rosette leaves. Indeed, accessions with higher GLK1 activity, arising from habitats with a high monthly variation in solar radiation levels, may show lower levels of photoinhibition at higher light intensities. Our results provide evidence for natural variation in GLK1 regulatory activities in vegetative leaves. This variation is associated with ecogeographic origin and can contribute to acclimation to high light conditions.

## Background

Photosynthetic activity is essential for the plant to capture energy from the sunlight and convert it to chemical energy needed for its activities, including growth and reproduction. At the same time, the plant also needs to control the imbalances between redox reactions created during photosynthesis, e.g. to avoid the damage produced by reactive oxygen species (ROS) [1–4]. In nature, these processes need to be coordinated under changing and fluctuating environmental conditions [5]. For example, northern European habitats have more dramatic seasonal changes in light and temperature conditions than Mediterranean climatic regions. Indeed, plants show a large variation in photosynthetic acclimation responses depending on their natural ecogeographic habitat range [5–7]. Although many studies have focused on how plants acclimate to a particular environment, little is known about the molecular mechanisms underlying the plasticity of acclimation responses to different environments.

Plants are able to acclimate photosynthetic activity to dynamic light conditions [4, 5], including under-using their photosynthetic capacity to avoid photooxidative stress [8]. They employ mechanisms that enable them to cope with excessive light or imbalance in light-dependent reactions and Calvin–Benson–Bassham (CBB) cycle, including thermal dissipation of excitation energy. Non photochemical quenching (NPQ) is for example associated with activity of the xanthophyll cycle and protonation of PSII antenna proteins [8]. Retrograde signaling pathways arising in the chloroplast evoke nuclear gene expression responses to acclimate photosynthetic activity and alleviate photooxidative stress [9, 10]. For example, reactive oxygen species (ROS) and cytosolic sugar levels have been proposed to act in retrograde signaling.

GOLDEN2-LIKE 1 (GLK1) and GLK2 TFs are main regulators of chloroplast biogenesis and photosynthetic activity in different flowering plant species [11, 12]. In Arabidopsis, GLK TFs promote the expression of many nuclear-encoded photosynthetic genes that are associated with chlorophyll biosynthesis and light-harvesting antenna proteins [13]. They bind and potentially regulate genes involved in acclimation to high light conditions [14, 15]. The *glk1 glk2* double mutant exhibits a reduction in NPQ [15], consistent with lowered Chl *b* levels.

The brassinosteroid (BR) pathway was shown to regulate GLK1 activity during chloroplast biogenesis. The BR-activated transcription factor BZR1 and its interaction partner PHYTOCHROME INTERACTING FACTOR 4 (PIF4) directly repress the expression of *GLK1* [16]. At post-translational level, the light-dependent GSK-3-like kinase BRASSINOSTEROID INSENSITIVE 2 (BIN2) phosphorylates GLK1, thereby promoting protein stability. At the same time, BIN2 phosphorylates the GLK1 repressor BZR1 resulting in reduced stability [17].

GLK1 target gene networks have been characterized previously by us and others [14, 18]. This offers an interesting resource and starting point to investigate natural variation in transcriptional regulatory networks underlying photosynthetic acclimation and its plasticity under different local habitat environments. Arabidopsis with its wide ecogeographical habitat range and genomic resource provides an excellent model system.

## RESULTS

### Gene expression diversity in the GLK1 regulatory network across *Arabidopsis* **accessions**

To understand the natural diversity and potential adaptation of the nuclear gene regulatory network controlling photosynthetic activity to different natural environments, we characterized the expression variation of GLK1 candidate target genes across 727 *Arabidopsis* accessions collected from a wide range of habitats. We previously performed chromatin immunoprecipitation followed by high throughput sequencing (ChIP-seq) for GLK1 [14]. As expected, those data show that GLK1 directly binds to genes with central roles in chlorophyll biosynthesis and photosynthesis activity, including genes needed for acclimation to high-light levels. To be conservative, we only used the top 100 GLK1 binding sites (FDR<7.2*10^-11^), and we defined a gene as a direct GLK1 target when its DNA-binding site resides within the region from 1 kb upstream to 1 kb downstream of the gene. This resulted in 136 candidate GLK1 target genes. To study their expression diversity across a large group of *Arabidopsis* accessions, we re-analyzed publicly available transcriptome data (RNA-seq) generated from rosettes of 727 Arabidopsis accessions collected just before bolting and grown under standard greenhouse long day conditions [19]. These accessions were collected from a global Arabidopsis habitat range (see **Suppl. Fig. 1a**). **Fig. 1a** shows the expression pattern of these 136 GLK1 potential target genes across the 727 accessions. We observed three main clusters of genes (**Fig. 1a**, Table S1): cluster G1 (n=68) was significantly enriched in photosynthesis and related Gene Ontology (GO) terms (**Suppl. Fig. 2**), G2 (n=63) showed a preponderance of genes involved in circadian rhythm (FDR<0.14, **Suppl. Fig. 2**). Cluster G3 contained only 5 genes and we did not test for GO enrichment due to low sample size. When testing for the enrichment of transcriptional regulators using ShinyGO v0.77 [20], G1 genes were enriched in targets of SQUAMOSA-BOX BINDING PROTEIN-LIKE 7 (SPL7) and Nuclear Transcription Factor Y SUBUNIT C2 (NFYC2) TFs (**Fig. 1a**). NFY TFs can interact with BBX TFs that have recently been identified as central regulators of photosynthetic gene expression [21, 22]. SPL7 is a regulator of copper signaling that is also required for ROS-detoxifying enzyme activities in plastids [23]. This indicates that G1 genes represent GLK1 target genes reflecting their function in chloroplast biogenesis and photosynthesis. Genes in cluster G2 were more enriched in CIRCADIAN CLOCK ASSOCIATED 1 (CCA1) and PHYTOCHROME INTERACTING FACTOR 1 (PIF1) target genes. The heatmap in **Fig. 1a** shows target genes of these TFs in the left. This suggests that cluster G2 represents potential target genes of GLK1 not directly related to photosynthetic activity. Indeed, GLK1 has been reported to also be involved in other functions such as anthocyanin biosynthesis and flowering [24, 25].

**Figure 1.**
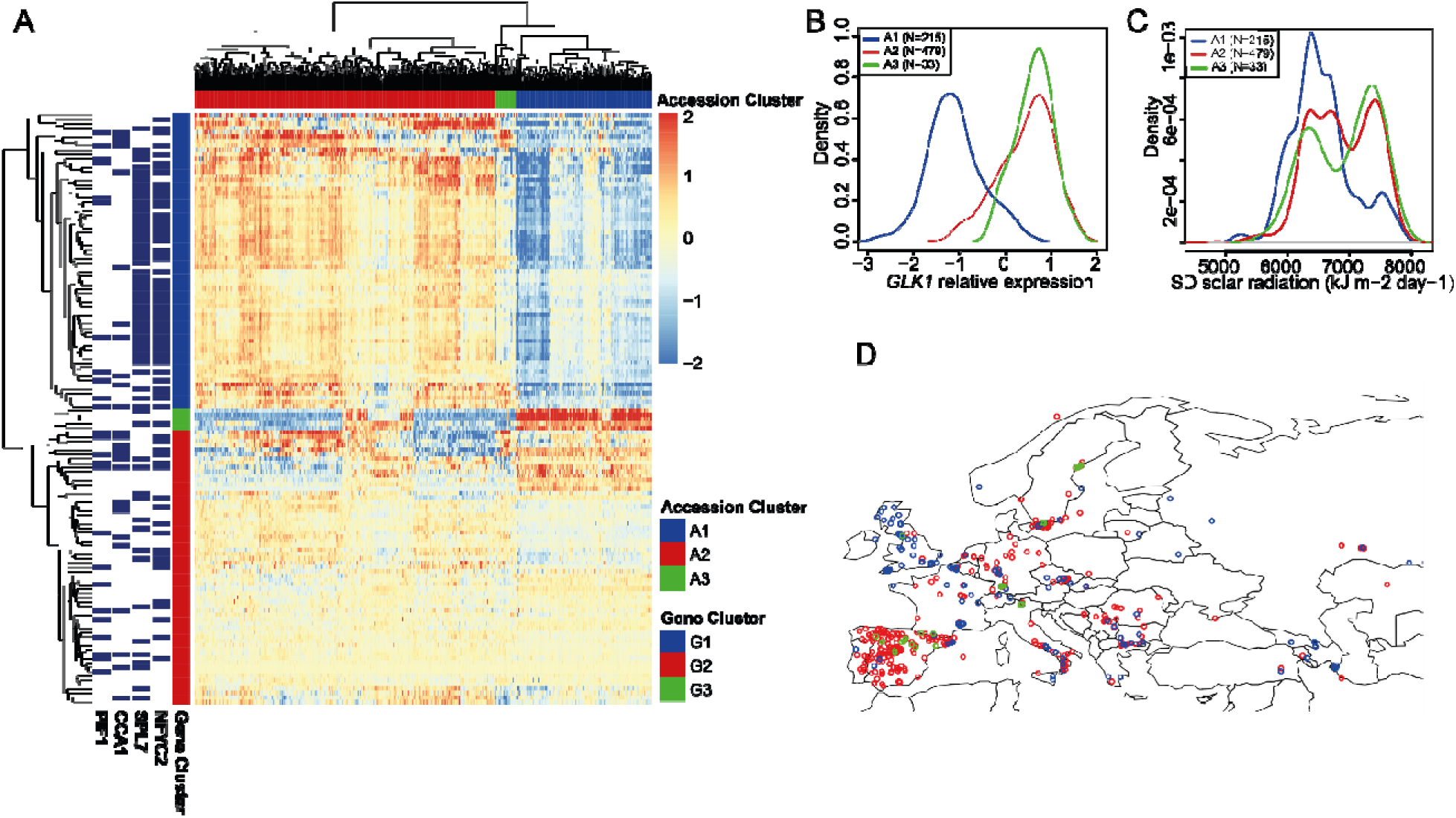
Gene expression diversity of GLK1 candidate target genes across 727 Arabidopsis accessions. **A)** Heatmap showing the relative gene expression of the candidate target genes (n=136) of the top 100 GLK1 binding events across 727 *Arabidopsis* accessions. Rows represent genes and column accessions. Three clusters of accessions and three clusters of genes were identified and labeled in the heatmap depending on their particular expression patterns. **B)** Expression of *GLK1* in the three clusters identified in Fig. 1A. The expression is significantly different (t-test; p-value<2*10^-16^) between A1 vs A2, and A1 vs A3, and slightly significant (p-value<0.03) between A2 vs A3. **C)** Distribution of the monthly standard deviation (SD) of solar radiation of the coordinates from the regions where the accessions were collected. Groups A1 and A2 were significantly different (p-value<10^-7^; t-test). **D)** Map representing the location of the 727 *Arabidopsis* accessions. Only a subsection of the map is plotted, see Suppl. Figure 1 for a complete map. A1 accessions are plotted in blue, A2 in red, and A3 in green.

Interestingly, the clustering algorithm identifies 3 different groups of *Arabidopsis* accessions with specific gene expression patterns (**Fig. 1a**, Table S2): accessions in group A1 (n=215) show a much lower expression of genes from group G1 (mainly photosynthetic genes) and of *GLK1* itself (**Fig. 1b**) compared to A2 (n=479) accessions. Meanwhile, candidate GLK1 target genes from the G2 group (non-photosynthetic functions of GLK1) did not show a strongly reduced expression like the G1 genes. Group A3 (n=33), which includes *Col*-0, was very similar in expression to A2 accessions, although with lower expression levels of *GLK2*. Genes of other known proteins affecting *GLK1* expression also show distinctive expression patterns in these accession groups (**Suppl. Fig. 3**): While *GENOMES UNCOUPLED 1* (*GUN1*) shows an expression pattern that is similar to *GLK1* in A1 vs. A2, the *GLK1* repressor *PIF4* shows a contrasting pattern. **Fig. 1d** shows the distribution of accessions in Europe, see **Suppl. Fig. 1a** for their global distribution. We focused our further analysis on clusters A1 and A2, since they show the most distinct gene expression profiles and contain most of the *Arabidopsis* accessions. The A1 accessions were collected from regions enriched in countries such as GBR and USA, while A2 accessions were enriched e.g. in Germany, Sweden, and especially Spain (**Supp. Fig. 1a**). This indicates that they may represent adaptation to different climatic environments. To further investigate this, we downloaded environmental variable values associated with the places of collection from WorldClim (version 2.0) [26]. Among the variables studied, we found that the inter-monthly standard variation of solar radiation (p-value < 1.1*10^-7^; t-test) was the most significantly different average between groups A1 and A2. Accessions from group A1 originate mainly from environments with lower inter-monthly variation in solar radiation (**Fig. 1c**). See **Table S3** for p-values of other variables studied.

Next, we aimed to characterize the general differences in gene expression patterns of the A1 and A2 accession groups. When analyzing the expression of all nuclear-encoded genes using DESeq2, we identified 1,179 genes as significantly differentially expressed in A1 compared to A2 accessions (FDR <0.01 and abs log2 FC>2; **Fig. 2a**). 824 of those genes were more active in accession group A1. On the other hand, 366 genes were more strongly expressed in the accession group A2. Together, these analyses suggest that the A1 and A2 accession groups show clear differences in genome-wide gene activities. To better characterize these genes, we performed GO enrichment analyses (**Fig. 2b-c**). Here, we found that genes related to reactive oxygen species (ROS) signaling are highly expressed in A1 (so linked with low *GLK1* expression), while genes involved in defense responses are more strongly expressed in accession group A2 (so associated with high *GLK1* activity).

**Figure 2.**
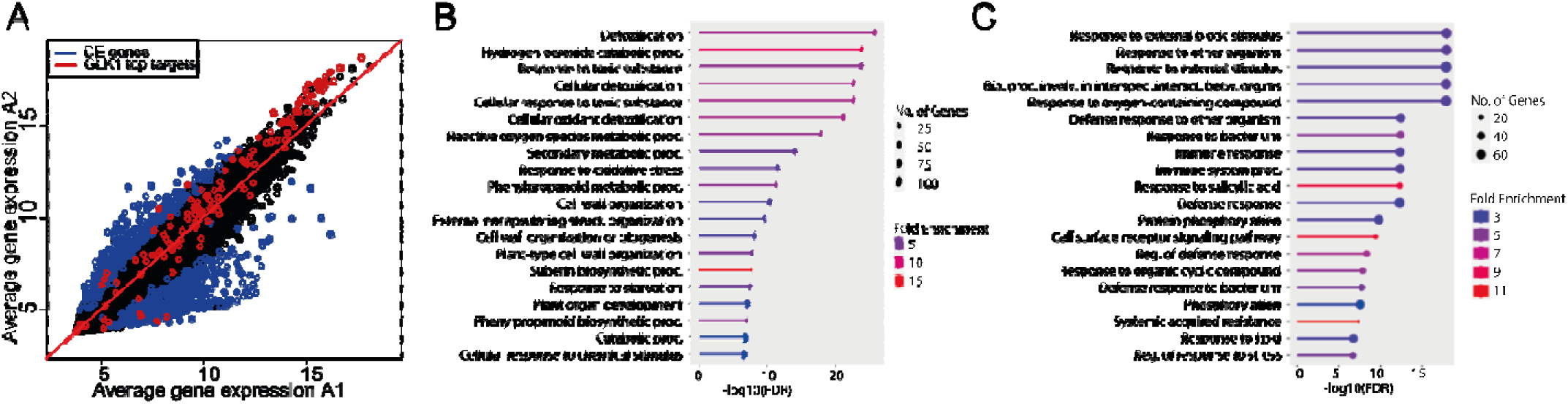
Differential gene expression signature in clusters A1 vs A2. **A)** Scatterplot showing the average expression of all genes in cluster A1 vs A2. In blue are indicated differentially expressed genes (FDR<0.01 & abs log2FC>2), and in red the target of GLK1 represented in Fig 1A. **B)** GO analysis (ShinyGO) of genes with a significantly increased expression in A1 vs A2. **C)** GO analysis (ShinyGO) of genes with a significantly increased expression in A2 vs A1.

Next, we studied the phylogenetic relationship among these accessions. The 1001 genomes consortium defined several admixture groups among the Arabidopsis accessions depending on their genome sequence similarity [27]. Indeed, we see that group A1 is enriched in the groups “Western Europe”, “Germany” and “Italy Balkan Caucasus”, while group A2 is enriched in the admixture groups “Central Europe”, “South-Sweden” and “Spain” (**Suppl. Fig 1c**), which indicates that different genetic structure is associated with the two accession groups.

In summary, we have identified at least two groups of *Arabidopsis* accessions with distinct transcriptome profiles related to *GLK1* activity. These two groups were collected from eco-geographic locations with different environmental properties, with solar radiation being the most statistically different (see Table S3 for the list and significance of all variables studied). These accessions represent different genetic structures, as they belong to different admixture groups.

### The genetic structure behind different *GLK1* activities across *Arabidopsis* accessions

Since we observed a different proportion of admixture groups in our *Arabidopsis* accession groups, we decided to characterize their genetic differences in more detail. We utilized the Single Nucleotide Polymorphisms (SNPs) and short insertions/deletions obtained from the 1001 Arabidopsis genomes project [27] to perform genome-wide association analysis (GWAS) to identify SNPs separating groups A1 and A2 (Chi-square test, PLINK software; **Suppl. Fig 4**). A large number of SNPs (n=77,424) were significantly associated with the trait at study (p-value<10^-10^; Table S4), indicating a complex genetic structure behind the phenotype studied. We filtered SNPs located outside exons or with a frequency lower than 25% in A1 or A2 accession groups. Among the remaining 6,918 SNPs, 115 SNPs were estimated by Variant Effect Prediction (VEP) to have a “HIGH” impact on their protein activity. Among them, 98 were predicted to introduce a new stop codon in the protein sequence. **Fig. 3** shows the genetic structure of the A1 and A2 groups, for SNPs predicted to have a “HIGH” or “MODERATE” impact on protein function.

**Fig 3.**
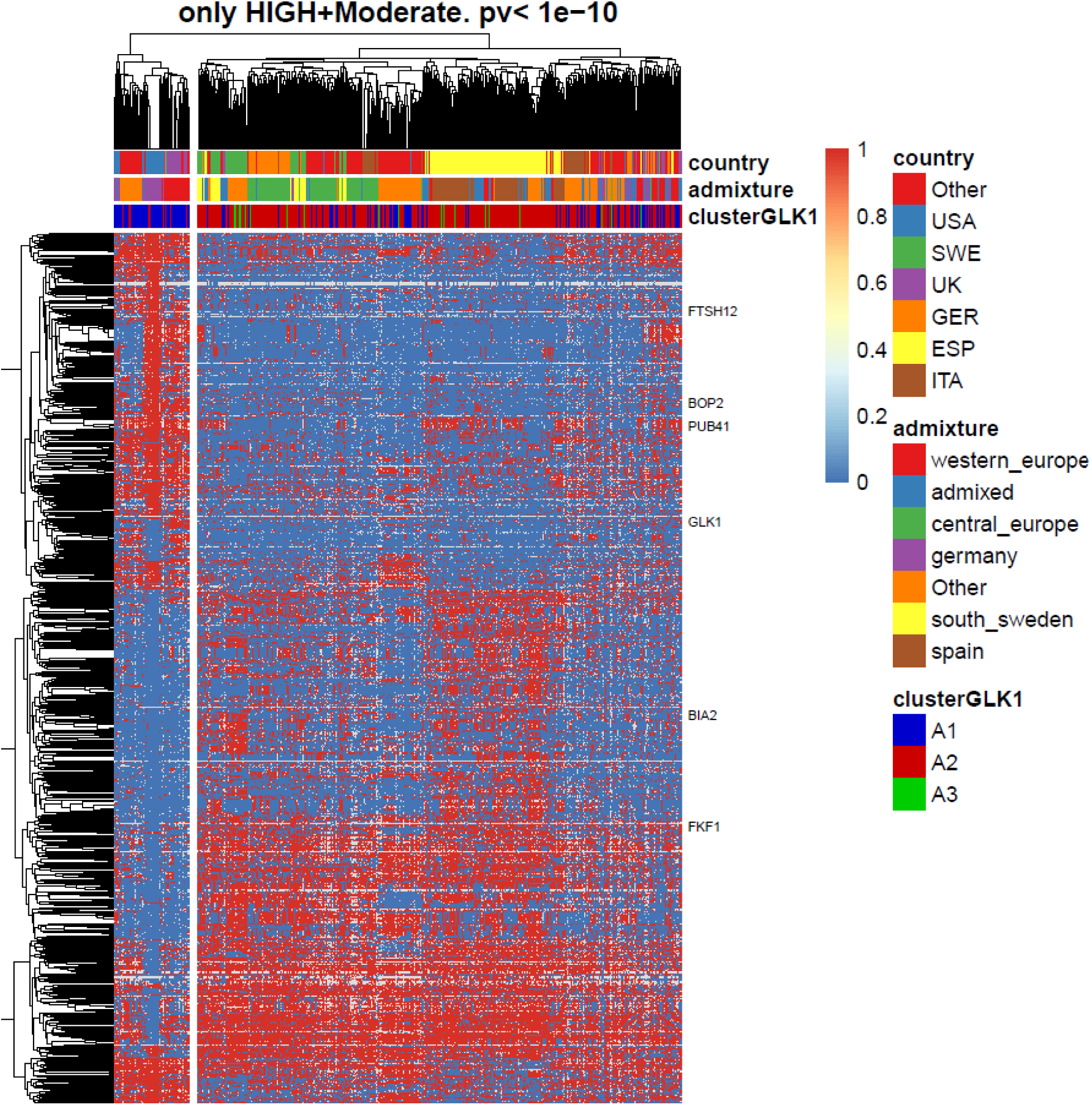
Genetic variation behind the different GLK1 activity. Heatmap showing SNPs among the Arabidopsis accessions studied. Only SNPs differently associated with A1 and A2 accession groups and with a predicted high or moderate impact in the gene sequence are shown. Genes predicted to have a high impact and mentioned in the manuscript are labeled on the right side.

Genetic variation with a “HIGH” impact on protein activity was found for genes that potentially could affect the GLK1 gene regulatory network. For example, FTSH PROTEASE 12 (FTSH12) serves as an import motor involved in the ATP-dependent translocation of pre-proteins through the TIC channel and is important for the retrograde/anterograde signaling pathways [28]. The mutation of this gene in Col-0 is reported to be embryo lethal [29]. However, many of our *Arabidopsis* accessions carry a premature stop codon without causing lethality which indicates a different gene-regulatory re-wiring that can compensate for this mutation. Another interesting example is *BRASSINOSTEROID INACTIVATOR 2* (*BIA2*), a gene that controls brassinosteroid homeostasis. Its overexpression is associated with change in expression of genes involved in photosynthesis (including *GLK1*), carbohydrate metabolism, and circadian rhythms [30]. Also affecting the brassinosteroid pathway, PLANT U-BOX 41 (PUB41) acts in the degradation of BZR1 [31]. Another gene affected encodes the NPR transcription factor BLADE-ON-PETIOLE 2 (BOP2) that is regulated by ROS signaling, controls boundary establishment in development and acts in plant defense [32]. Other genes harboring mutations in subsets of the accessions are e.g. *GLK1* itself and *STROMAL ASCORBATE PEROXIDASE* (*SAPX*) that mediates ROS scavenging in the chloroplast stroma [33].

In summary, we identified a complex genetic structure behind these two distinct clusters of Arabidopsis accessions. Focusing on SNPs predicted to cause a change in protein activity (i.e. by introducing premature stop codons), we determined candidate genes that may contribute to natural variation in the levels of GLK1 activity and thereby differential regulation of its target networks.

### Gene-regulatory network analysis

To better understand the differences in gene activities observed between A1 and A2 accession groups, we predicted the gene regulatory network (GRN). Based on the analysis of genetic polymorphisms, we focused on genes involved in GO categories related to chlorophyll metabolism, brassinosteroid signaling and retrograde signaling (see Methods; n=216 genes) using GeneNet [34] for the group of A1 and A2 accessions independently. We did not preselect for TFs when running GeneNet as many regulators of the chlorophyll metabolism are not only TFs, but also e.g. transporters and posttranscriptional or posttranslational regulators, such as RNA binding proteins. GeneNet infers large-scale gene association networks using graphical Gaussian models that represent dependencies among genes calculated based on partial correlations. Partial correlations measure the dependence of two variables given a set of confounding variables (other genes in this context) and therefore provide a closer measure of direct effects.

We initially used the GRN of A2 as a reference as it is more similar at the *GLK1* target expression level to Col-0 compared to A1 (Fig 1a). Among the 216 genes considered, 1,394 interactions were found significant in the A2 accessions (p-value<0.05; Table S5). To predict genes regulating *GLK1* or genes acting in the same regulatory module, we selected the top 36 gene interactions in the neighborhood of *GLK1* predicted by GENENET (p-value< 0.01), and plotted these interactions in a graphical network (**Fig. 4**). GLK1 was predicted to be associated with (in order of importance) GLK2, LZF1, GNC, BEH1, and BCM1. Indeed, GLK2 may cross-regulate with GLK1 as they bind each other’s promoters [14, 18]. LZF1/BBX22 is a B-box protein involved in the positive regulation of photomorphogenesis [35]. Other B-box TFs are regulated by GLK1 [14, 25]. GNC and GLK1 have overlapping functions coordinating chloroplast development [15]. BEH1 is a TF homolog of BEZ1 controlling brassinosteroid signaling and photosynthetic activity [36]. Lastly, *BCM1* encodes a Mg-dechelatase that catalyzes Chlorophyll *a* degradation and is regulated by GLK1 [37].

**Figure 4.**
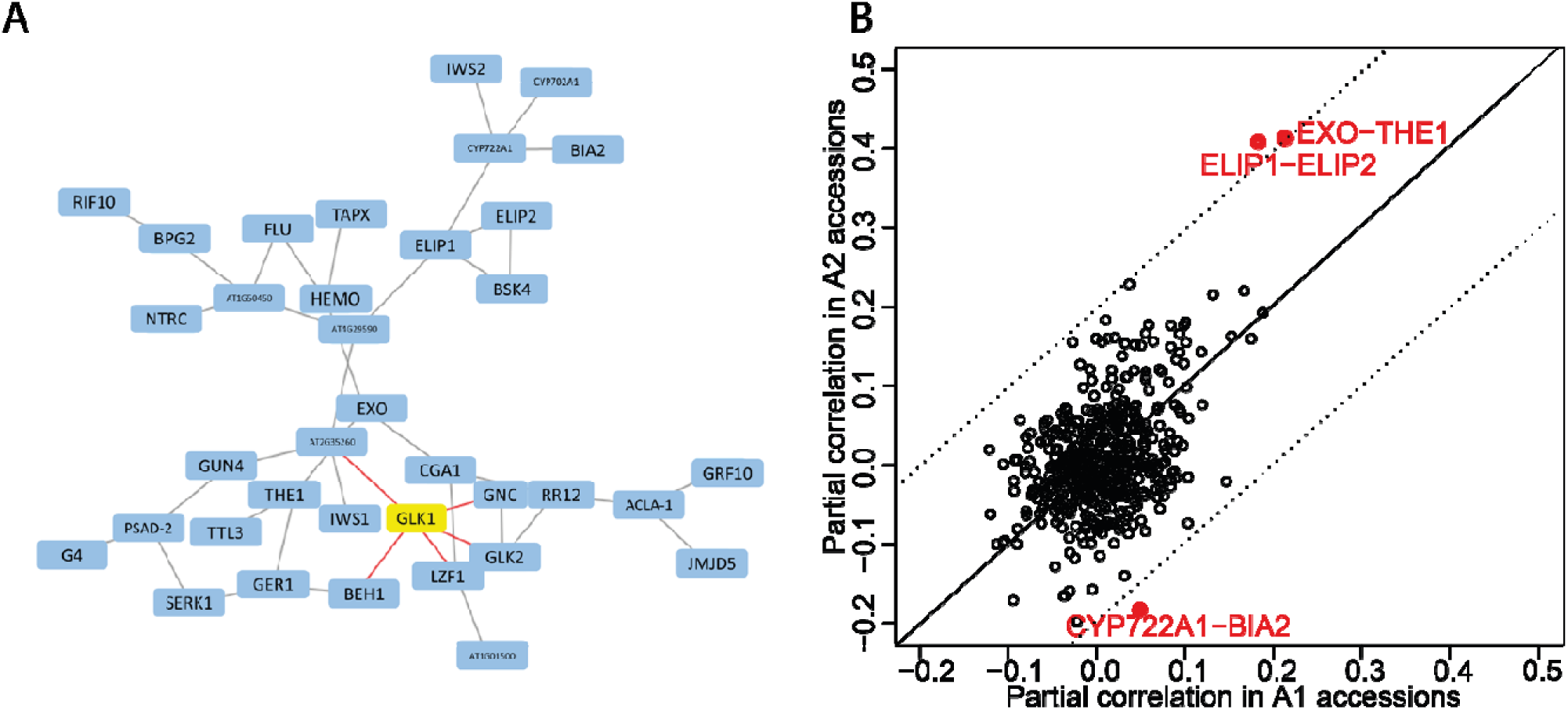
Gene regulatory network differences. **A** GRN around *GLK1* in the A2 group of accessions. **B** Interaction strength in the predicted in the A2 accession (y-axis) versus A1 accession (x-axis) for all possible interactions of the genes depicted in **A**.

In a next step, we determined the GRN for the set of A1 accessions (Table S6). In order to identify the main differences in the interactions of these two networks, we plotted the interaction strength predicted by GENENET in the A1 vs. A2 GRNs, for all interactions found significant (p-value<0.01) in the A2 GRN. Among all the interactions, the links between BIA2-CYP722A1, EXO-THE1 and ELIP1-ELIP2 were the most different. For the BIA2-CYP722A1 interaction, this could be explained by the fact that our GWAS analysis detected an SNP causing a new stop codon in BIA2 with different frequencies in the A1 and A2 accessions. The cell-wall localized protein EXORDIUM (EXO) and the receptor kinase THESEUS1 (THE1) control cell growth and act in the brassinosteroid pathway [38, 39]. ELIP1 and ELIP2 are two chlorophyll a/b binding proteins that have been suggested to play roles in protection from photo-oxidative stress [40], and thus may contribute to environmental acclimation pathways converging on *GLK1* activity. In summary, by comparing the gene regulatory network centered on *GLK1* activity in A1 vs A2, we revealed multiple links with specific genes in the brassinosteroid pathway, with a strong candidate gene affecting the natural variation in the levels of *GLK1* activity being *BIA2*. Indeed, previously it was found that *BIA2* overexpression (BIA2-OE) leads to elevated *GLK1* activity (FDR<4*10^-22) and *GLK2* (FDR<1.36*10^g-4) [30]. Additionally, we found that the *bia2* loss-of function alleles in natural accessions are correlated with low *GLK1* activity (**Suppl. Fig. 5a**). A comparison of transcriptome levels in accession cluster A1 and A2 showed that accessions with high expression of *BIA2* (A1) show a BIA2-OE transcriptome signature (**Suppl. Fig. 5b**).

### *GLK1* expression variation is linked with fluctuations in solar radiation

Given the clear difference in genetic structure and photosynthesis-related expression patterns between the two groups of Arabidopsis accessions studied here, and their association with different natural environments, we speculated that these genetic and expression changes may represent an adaptation to different environmental conditions. Indeed, these two subpopulations are significantly associated with different distributions of inter-monthly solar radiation variation.

The average solar radiation in a particular month depends on the day’s length and the solar intensity during the day. In this way, countries with a variable day length (e.g. Sweden), or with a variable solar intensity across the year (e.g. Spain) have a more variable solar radiation than other countries, such as the GBR. Indeed, long or short-day conditions are reported to impact the gene activities associated with photosynthetic pathways [41]. Also, excess light can result in the production of ROS, and plants have evolved the mechanism of non-photochemical quenching (NPQ) to dissipate excess light energy as heat to protect the photosynthetic machinery (for review, see [42]). Rungrat et al. showed that Arabidopsis adapted to environments with different solar and temperature levels by adjusting their NPQ activity [43]. They profiled 284 Arabidopsis accessions in two sets of conditions reflecting seasonal and regional (coastal vs. inland) climates, by simulating specific gradients in light intensity and temperature. The maximum light intensity was 150 µmol m^-2^ s^-1^ (coastal) or 300 µmol m^-2^ s^-1^ (inland), and photosynthetic parameters were measured by pulse amplitude modulation (PAM) fluorometry. Among the profiled accessions, 12 belong to our group A1, and 32 to the A2 group. Comparing photosynthetic parameters across conditions and accession groups (**Fig. 5a**) revealed that maximum quantum efficiency of photosystem II as measured by dark-adapted Fv/Fm (QY-max) is higher under inland conditions than under coastal conditions for accession group A1 (t-test, p-value<0.010). In contrast, QY-max of A2 accessions is not significantly different between inland and coastal conditions (p-value<0.543) (**Fig. 5b**). In line with this, A2 accessions display a significant increase in the photoprotection parameter NPQ_Lss (steady state non-photochemical quenching) (t-test, p-value<2.6*10-5, **Fig. 5c**), comparing “coastal” to “inland” conditions. There are no significant differences in NPQ_Lss for the A1 accessions when comparing these two environmental conditions (p-value<0.5). It is tempting to speculate that the A2 accessions, natural from countries like Spain and Sweden with high variation in solar radiation levels, show a stronger adaptive capacity in protection against photoinhibition, while photosynthetic efficiency (QY_Lss) is similar in A1 and A2 (**Fig. 5d**). In contrast to that, the A1 accessions, natural from low variation solar radiation environments like GBR, may not have the capacity to modulate protection against photoinhibition in adaptation to seasonal light differences.

**Figure 5.**
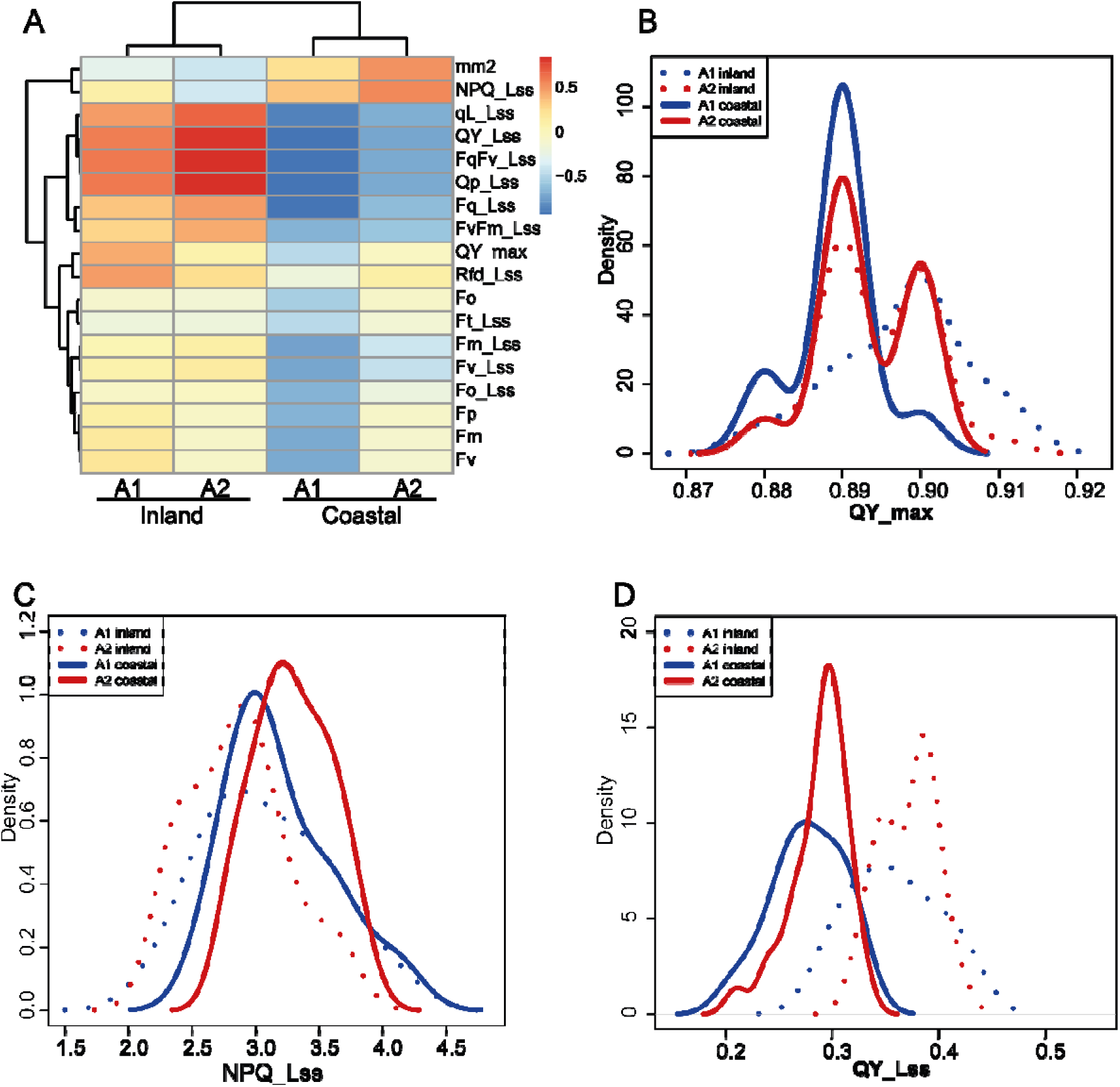
Photosynthetic activity measures across Arabidopsis accessions. The two groups of accessions studied have different photosynthetic activity. **A** Average photosynthetic value (column) among the accession groups and environmental conditions considered (rows). **B** Distributions for QY_max, the distributions for A2 accessions are not significantly different between the conditions studied (t-test, p-value<0.543). A1 accessions are significantly different when growing in the two environments (t-test, p-value<0.010). **C** Distributions for NPQ_Lss, the distributions for A1 accessions are not significantly different between the conditions studied, however, A2 accessions are significantly different when growing in different environments considered (t-test, p-value<2.6*10^-5^). **D** Distributions for the parameter QY_Lss in the A1 and A2 accession groups for the two different sets of conditions.

### Discussion

The acclimation of photosynthetic activities to variable environmental regimes requires regulatory adjustments at post-translational, post-transcriptional and transcriptional levels, and the coordination of gene activities in both plastid and nuclear genomes. GLK transcription factors function not only as central regulators of chloroplast biogenesis, but have also been found to act in response to abiotic and biotic environmental factors. Here, we showed that the levels of GLK1 activity in rosette leaves vary across natural Arabidopsis accessions, and that this variation is linked with eco-geographic environmental factors, especially seasonal fluctuations in solar radiation levels. GWAS and gene-regulatory network analyses identified brassinosteroids and ROS signaling as potential upstream pathways modulating *GLK1* activity in the different accession groups.

Acclimation to varying light conditions is mediated by a combination of short-term metabolic shifts and posttranslational regulation of the photosynthetic apparatus. Prolonged exposure to high-light conditions is mediated by additional adjustment in the photosynthetic status, e.g. mediated by changes in enzyme and antenna protein composition, and stomata aperture [44]. This in turn results in changes in the maximum photosynthetic rate (Pmax). These acclimation responses require retrograde signaling from the plastids to the nucleus. Different types of retrograde signals mediating plant acclimation to environmental conditions have been proposed, including ROS signaling and cytosolic sugar levels [10]. *GLK1* is a target of GUN1-mediated retrograde signaling [45]. This has been suggested to help the plants protect against photo-oxidative damage. At the post-translational level, GLK1 was found to be regulated by retrograde signals that influence protein stability [46]. More recently, it was shown that BIN2-mediated phosphorylation of GLK1 promotes chloroplast biogenesis by enhancing GLK1 protein stability and activity in the light [17]. This complexity of transcriptional and posttranscriptional regulation makes GLK1 a candidate factor for mediating acclimation of photosynthetic activity to different habitats. Our findings suggest that *GLK1* activity is genetically variable across Arabidopsis accessions, and that the variation is mediated by variants in different genes that contribute to finetuning GLK1 expression. These genes include known regulatory pathways of GLK1, for example brassinosteroid signaling genes such as *BIA2*, a regulator of brassinosteroid homeostasis [30]. Another factor that is associated with variation in GLK1 activity is BLADE-ON-PETIOLE 1 (BOP1), which was also shown to be regulated by brassinosteroids. Interestingly, BOP1 is a TGACG sequence-specific binding protein (TGA) TF and shows conserved protein residues that are known to be regulated in other TGA TFs by the intracellular redox status [47], thus providing a potential link with ROS signaling. The high number of genetic variants potentially contributing to variation in GLK1 gene activities in rosette leaves across the studied accessions suggest frequent and evolutionary dynamic adaptation of this regulatory system to local habitats. Indeed, we identified a different ecogeographic distribution of accessions with low vs. high *GLK1* levels. The complexity of genetic variations suggests multiple independent origins of genetic variation contributing to the diversity of GLK1 upstream pathways. To which extent this is linked with coupled genetic variation in the downstream pathways of *GLK1* will be an interesting question for future research.

## CONCLUSIONS

In summary, our findings suggest that variation in GLK1 regulatory pathways, such as the brassinosteroid pathway, mediates differences in the regulation of basal *GLK1* levels in different Arabidopsis accessions. Together with variation in downstream networks, this results in different adaptive potential to environmental conditions, in particular fluctuating light activities, by elevating or lowering the responsiveness of GLK1 target genes. Future experiments should test the roles of the identified candidate TFs and pathways, in particular brassinosteroid and ROS signaling in the regulation of GLK1 responsiveness in different natural environments. This will also shed light on the interplay between biogenic and operational retrograde signaling mechanisms converging on this important transcription factor.

## METHODS

### RNA-seq data analysis

Fastq files from publicly available RNA-seq data were downloaded from Sequence Read Archive (SRA; https://www.ncbi.nlm.nih.gov/sra) with ID PRJNA319904. Individual fastq files were trimmed with Trimmomatic v0.36 (default parameters), and mapped to the TAIR9 Arabidopsis genome using STAR v2.7.2b with options *--outFilterMultimapNmax 2 -- outMultimapperOrder Random --alignIntronMax 3000 --outSAMstrandField intronMotif -- outFilterIntronMotifs RemoveNoncanonical.* Mapped reads were assigned to genes (intron/exon) using *featureCounts* from the package *subread v1.6.4* with parameters *-s 2 -p -C - M -t gene -g gene_id*. Non protein-encoding genes or genes encoded in the organelle genomes were filtered out at this step. Next, individual sequencing libraries with less than 10^6^ mapped reads were filtered out. The remaining libraries were combined depending on the *Arabidopsis* accession used. This resulted in raw read counts for 727 Arabidopsis accessions and 27,445 genes. Next, the data was normalized using the function *varianceStabilizingTransformation* from the R package *Deseq2 v1.34.0.* When needed relative expression per gene was calculated as the expression of a particular gene in one particular *Arabidopsis* accession minus the average expression of that gene across all accessions.

The function *pheatmap* from the R package *pheatmap* v1.0.12 was used to cluster the relative expression of the candidate GLK1 targets across all Arabidopsis accessions studied. For this, the top 100 GLK1 binding sites were obtained from [14]. Candidate GLK1 targets were defined as protein-coding genes with a top 100 GLK1 binding sites in the 1kb upstream, inside, and 1kb downstream region of the gene. This resulted in 136 genes.

### Climate data

Average monthly climate data were downloaded from the WorldClim v2.0 database using a spatial resolution of 30s (∼ 1 km^2^) across the years 1970-2000 [26]. The R package RGDAL v1.6-7 was used to obtain the climate data from the location where the different *Arabidopsis* accession were collected as reported by the 1001Genome project. The average climate values were obtained by calculating the average value across the 12 months of the year, and the standard variation was obtained by calculating the standard deviation across the 12 months of the year.

### GWAS analysis

SNP data for the *Arabidopsis* accession studied were downloaded from the 1001 Genome project v3.1, file name: 1001genomes_snp-short-indel_only_ACGTN.vcf.gz which only contains genetic variants located in the nuclear genome. GWAS analysis (Chi-square test for the A1 versus A2 groups) was performed with the *Plink* software v1.90b6.21, with parameters: *---assoc-allow-no-sex --pfilter 0.05.* In addition, only variants with a p-value lower than 10^-10^, with a frequency bigger than 25% in the A1 or A2 *Arabidopsis* accession groups as estimated by Plink were retained for further analysis, and plots.

### Gene regulatory network reconstruction

Gene expression profiles were obtained using the analysis described before. Only genes belonging to the next gene ontologies and their children were used: “chlorophyll metabolic process” (GO:0015994), “chloroplast-nucleus signaling” (GO:0010019), “brassinosteroid homeostasis” (GO:0010268) and “brassinosteroid mediated signaling pathway” (GO:0009742). The list of genes belonging to these ontologies was downloaded from the TAIR website (https://www.arabidopsis.org). This results in 214 genes. Gene regulatory networks were estimated for the A1 and A2 accession groups independently using the function *ggm.estimate.pcor* of the R package *GeneNet* with parameters method=”dynamic”. Only edges with a p-value associated lower than 0.05 were considered. To predict *GLK1* regulatory network (Fig. 4), we selected genes in the *GLK1* network as the genes with a p-value<0.01 to be associated with *GLK1*. To provide more depth to the network, genes directly associated with the previously selected GLK1-associated network were also selected. This was done 5 steps upstream of GLK1, which resulted in 36 genes. For these genes, the association was plotted in Cytoscape v3.10.

### Re-analysis of photosynthesis activity

Photosynthesis activity data for 271 Arabidopsis ecotypes and genotypes were downloaded from the supplementary material of [43]. Only data from ‘leaf 14’ were used. Among their profiled accessions, 12 belong to our group A1, and 32 to the A2 group of *Arabidopsis* accessions.

## DECLARATIONS

## Abbreviations

TF: transcription factor
GRN: gene regulatory network
GWAS: genome-wide association mapping

## Ethics approval and consent to participate

Not applicable

## Availability of data and materials

Datasets and scripts are available upon request

## Consent for publication

Not applicable

## Competing Interests

The authors declare no competing interests.

## Funding

K.K. wishes to thank the DFG for funding (project no. 270050988 (C05); 458750707). T.K. wishes to thank the DFG for funding (project no. 270050988 (C01)). J.M. wishes to thank the DFG for funding (project no. 407463262).

## Authors’ contributions

J.M. performed the data analyses. J.M. and K.K. wrote the main manuscript text and prepared the figures. C.G. and T.K. edited the manuscript. All authors reviewed the manuscript.

## Acknowledgements

The authors wish to thank Christian Schmitz-Linneweber for valuable comments on an earlier version of the manuscript.

## SUPPLEMENT

### Supplementary Figures

**Suppl. Fig 1.**
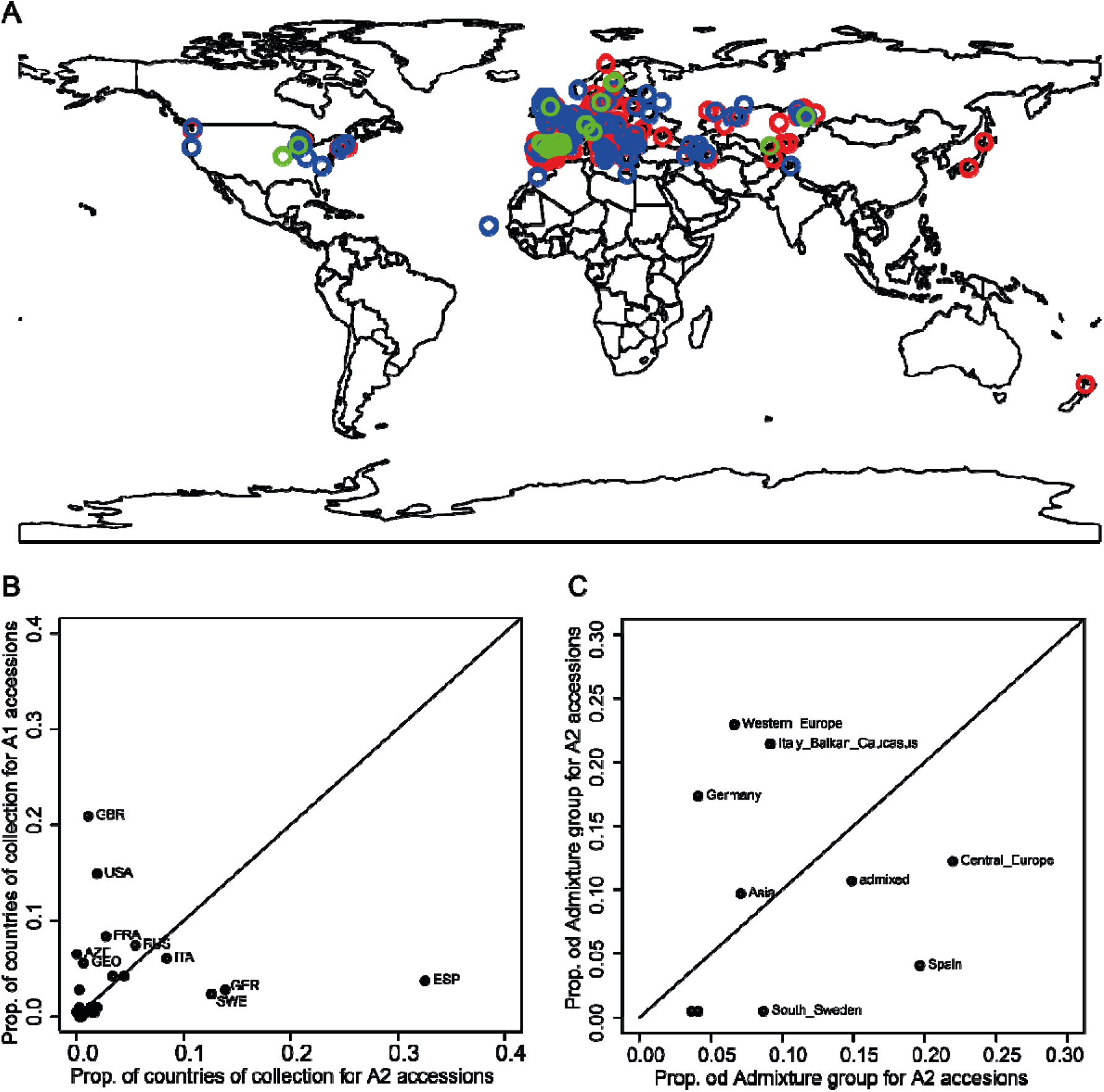
Spatial distribution of the studied *Arabidopsis* accessions. **A)** World map plotting the location from where accessions were collected. A1 accessions are plotted in blue, A2 in red, and A3 in green. **B)** Proportion of country of origin is plotted for accessions belonging to clusters A1 and A2. **C)** Proportion of admixture groups as identified by the 1001 Genomes project [27].

**Suppl. Fig 2.**
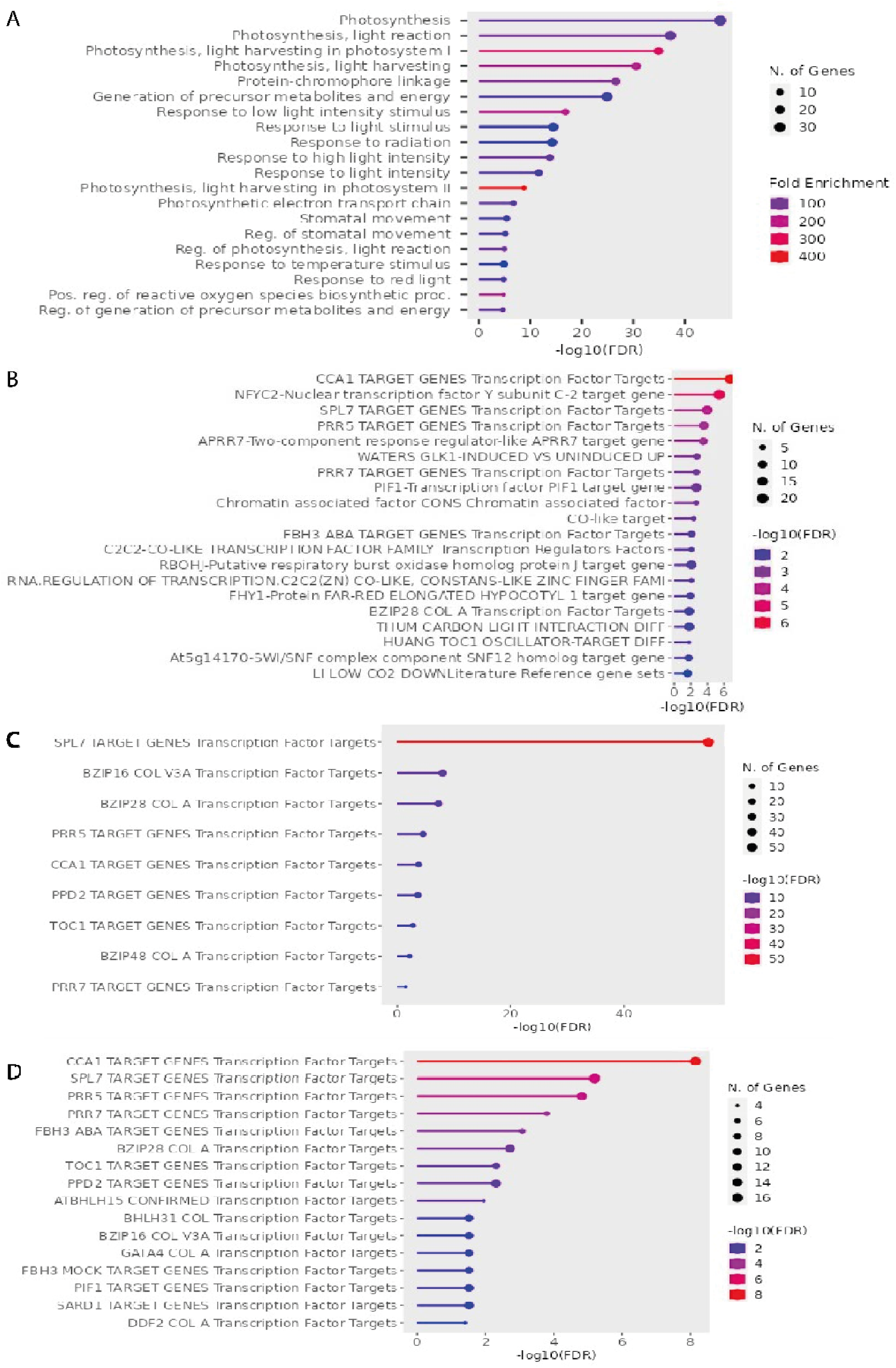
GO analysis of DE genes among targets of the top 100 GLK1 BSs. GO analysis was performed with ShinyGO v0.77. The GO enrichment for G1 genes is shown in **A**, and for G2 genes is shown in **B**. ShinyGO can also test the enrichment among the Plant Gene Set Annotation database [48]. The enrichment for G1 genes is shown in **C** and for G2 genes in **D**.

**Suppl. Figure 3.**
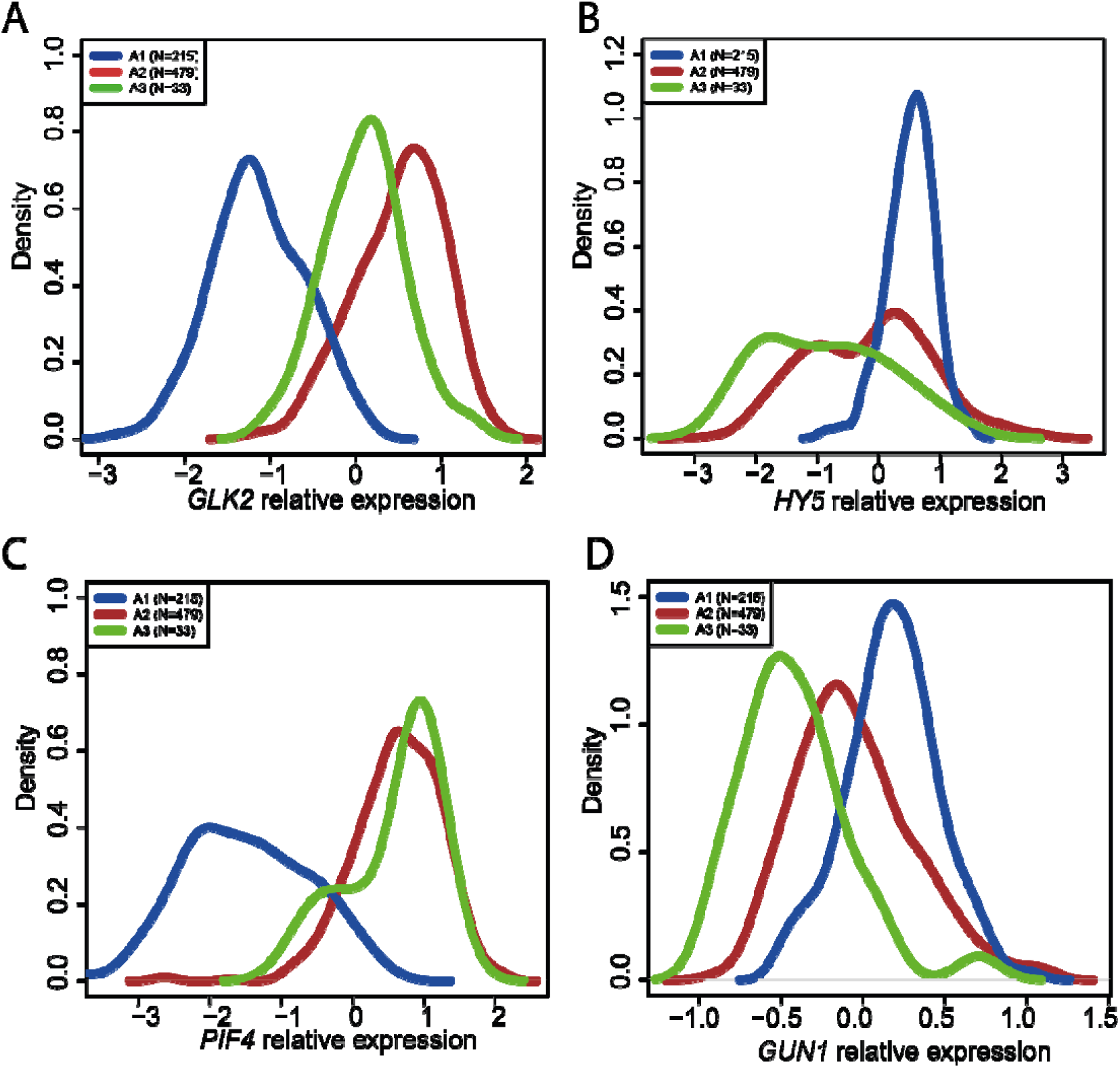
Expression of known *GLK1* regulators in different *Arabidopsis* accession groups. The expression distribution of *GLK2* (A), *HY5* (B), *PIF4* (C) and *GUN1* (D) is shown for each group of accessions defined in Fig. 1a.

**Suppl. Figure 4.**
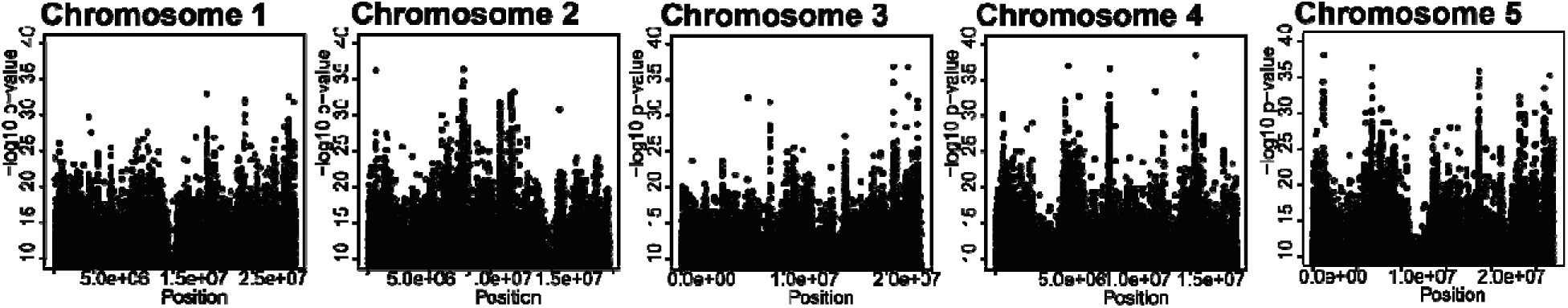
Distribution of -log 10 p-values obtained by PLINK when testing (Chi-square test) for association between the group of accessions A1 versus A2 for each chromosome. Typically, a fixed p-value threshold of 5*-8 is widely used to identify association between genetic variants and a trait of interest [49].

**Suppl. Figure 5.**
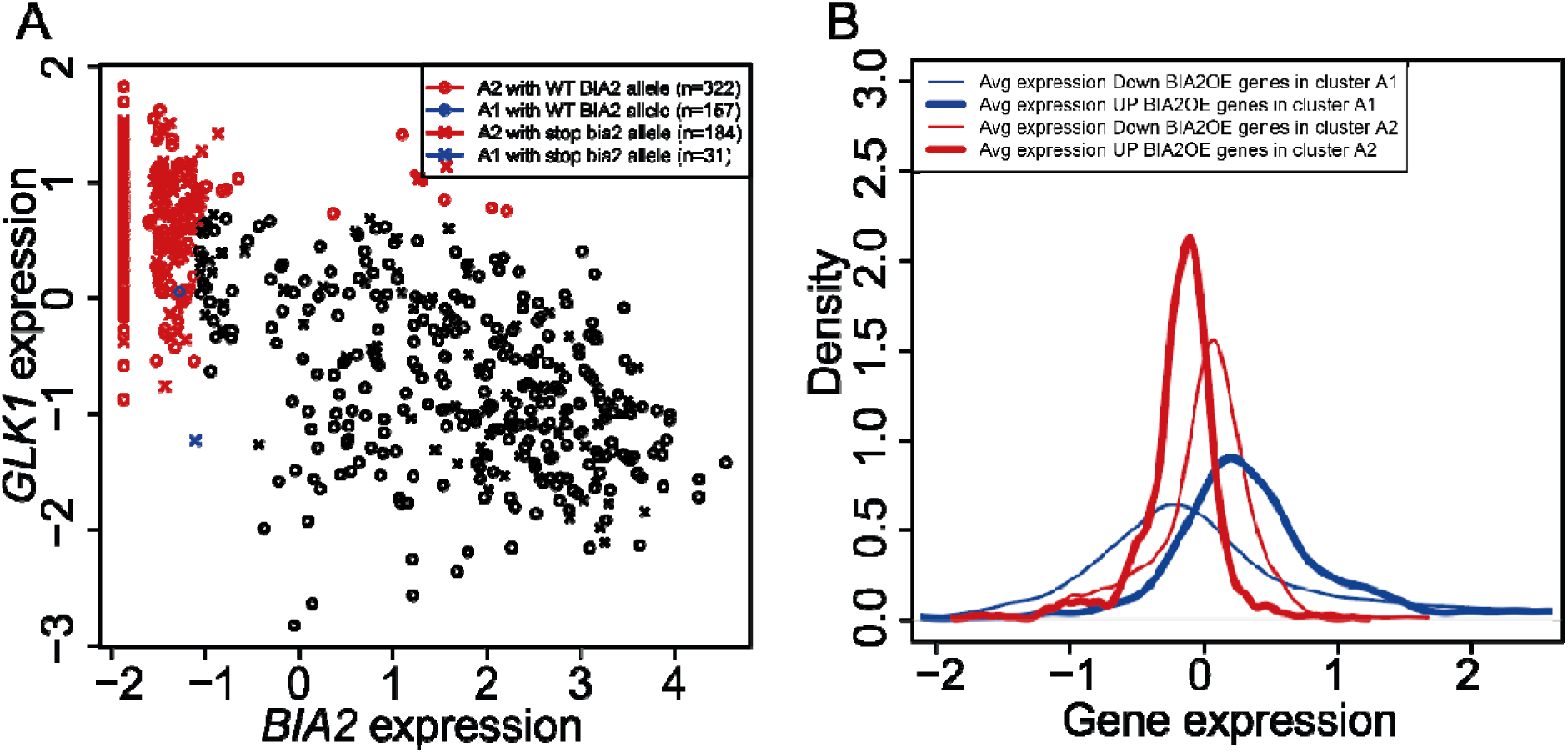
**A** Relation of *GLK1* and *BIA2* expression, as well as impact of mutation causing stop codon in the *BIA2* gene. **B** Expression signature of accessions with high *BIA2* expression fits with the observed consequence of *BIA2* overexpression in *Col*-0.

## Notes

### Competing Interest Statement

The authors have declared no competing interest.

